# MG132 induces progerin clearance and improves disease phenotypes in fibroblasts of patients affected with Hutchinson-Gilford Progeria-like syndromes

**DOI:** 10.1101/2021.04.14.439612

**Authors:** Karim Harhouri, Pierre Cau, Frank Casey, Guedenon Koffi Mawuse, Yassamine Doubaj, Lionel Van Maldergem, Gerardo Mejia-Baltodano, Catherine Bartoli, Annachiara De Sandre-Giovannoli, Nicolas Lévy

**Affiliations:** Aix Marseille Univ, INSERM, U 1251-MMG, Marseille, France; Progelife, Marseille, France; Royal Belfast, Pediatric Cardiology, Hospital for Sick Children, Belfast BT9 7AB Northern Ireland; CHU Sylvanus Olympio de Lomé, Unité de Génétique Humaine, Lomé BP 1515, Togo; Département de Génétique Médicale, Institut National d’Hygiène, 11400 Rabat, Morocco; Centre de génétique humaine, CHU Université de Franche-Comté, Besançon, France; Departamento de Genética Ministerio de Salud de Nicaragua; Hospital Infantil “Manuel de Jesús Rivera”, Managua, Nicaragua; AP-HM, Hôpital d’Enfants de la Timone, Département de Génétique Médicale, Marseille, France; Biological Resource Center (CRB-TAC), Assistance Publique Hôpitaux de Marseille, La Timone Children’s Hospital, Marseille, France

**Keywords:** Progeria-like, MAD-B, Progerin, Prelamin A Δ90, Prelamin A Δ35, MG132, autophagy, inflammation

## Abstract

Progeroid Syndromes (PS), including Hutchinson-Gilford Progeria Syndrome (HGPS, OMIM #176670), are premature and accelerated aging that clinically resemble some aspects of advancing physiological aging. Most classical HGPS patients carry a *de novo* point mutation within exon 11 of the *LMNA* gene encoding A-type Lamins. This mutation activates a cryptic splice site leading to the deletion of 50 amino acids at its carboxy-terminal domain, resulting in a truncated and permanently farnesylated Prelamin A called Prelamin A Δ50 or Progerin that accumulates in HGPS cell nuclei and is a hallmark of the disease. Some patients with PS carry other *LMNA* mutations affecting exon 11 splicing, leading to defects in nuclear A-type Lamins and are named “HGPS-like” patients. They also produce Progerin and/or other truncated Prelamin A isoforms (Δ35 and Δ90). We recently found that MG132, a proteasome inhibitor, induced progerin clearance in classical HGPS through autophagy activation and splicing regulation. Here, we show that MG132 induces aberrant prelamin A clearance and improves cellular phenotypes in HGPS-like patient cells. These results provide preclinical proof of principle for the use of a promising class of molecules toward a potential therapy for children with HGPS-like, who may therefore be eligible for inclusion in a therapeutic trial based on this approach, together with classical HGPS patients.

## INTRODUCTION

Progeroid syndromes (PS) are a group of very rare genetic disorders associated with clinical features that mimic physiological ageing. Hutchinson-Gilford Progeria Syndrome (HGPS) is the most prevalent and widely studied syndrome among PS. Estimates indicate that the prevalence of HGPS is approximately one in 4 million children (Hennekam, 2006). HGPS is characterized by premature and accelerated aging with rapidly growth retardation, thin skin, loss of subcutaneous fat, alopecia, osteoporosis, and cardiovascular disease. HGPS patient’s death occurs at the mean age of 14.6 years (Gordon et al., 2014), almost exclusively due to heart attack or stroke caused by atherosclerosis. In 2003, we and an US group identified a recurrent de novo point mutation (c.1824C>T, p.G608G) in the *LMNA* gene (1q21) encoding A-type lamins as the most frequent cause of classical progeria (De Sandre-Giovannoli et al., 2003;Eriksson et al., 2003). In physiological conditions, *LMNA* encodes lamins A and C through alternative pre-mRNA splicing. Lamin A/C are major components of the nuclear Lamina, a protein meshwork located underneath the inner membrane of the nuclear envelope and dispersed through nuclear matrix [4]. The HGPS mutation activates a cryptic splice site in prelamin A-encoding mRNAs, mainly regulated by the serine–arginine rich splicing factor 1 (SRSF-1) (Lopez-Mejia et al., 2011), leading to the production of a truncated and permanently farnesylated prelamin A precursor (called progerin). Progerin cannot be properly post-translationally processed to mature lamin A and thus accumulates at the cell nuclear periphery. Progerin exerts a series of toxic, dose-dependent, dominant negative effects including altered heterochromatin dynamics, DNA damage repair defects, chronic inflammation, proliferation slowdown and accelerated senescence (Gonzalo, Kreienkamp, & Askjaer, 2017). Progerin intranuclear accumulation has thus been identified as a major HGPS pathophysiological target, and is being involved in most, if not all, of the 9 hallmarks of physiological aging (Lopez-Otin,Blasco, Partridge, Serrano, & Kroemer, 2013).

A wide spectrum of treatment strategies with different specificities, targeting several processes, has been proposed to correct the defects in HGPS: (i) to “repair” the disease-causing mutation; (ii) to block pre-mRNA aberrant splicing leading to progerin mRNA production; (iii) to reduce the toxicity of isoprenylated and methylated progerin; (iv) to induce progerin clearance; (v) to decrease the noxious downstream effects linked to progerin accumulation (Cau et al., 2014;Harhouri et al., 2018). However, targeting only one pathophysiological event of progeria and related diseases would not result in a reversal of the pathological phenotypes of such segmental disorders that affect multiple tissues, hence therapeutic approaches targeting several mechanisms triggering the disease are needed, that can be envisaged for all related syndromes having common pathophysiological mechanisms. These approaches should succeed in lowering amount of aberrant prelamin A isoforms at different levels, including their decreased production, increased degradation as well as counteracting downstream toxic effects. We demonstrated that proteasome inhibitor, MG132, not yet FDA/EMA-approved, induce a spectacular progerin inhibition through a dual action: MG132 induces progerin degradation through macroautophagy and strongly reduces progerin production through downregulation of SRSF-1 and SRSF-5, controlling prelamin A mRNA aberrant splicing. MG132 treatment improves cellular HGPS phenotypes *in vitro* and injection of the drug in skeletal muscle of a mouse model of progeria (*Lmna*^G609G/G609G^) locally reduces SRSF-1 expression and progerin levels (Harhouri et al., 2017).

Besides typical HGPS, there are other forms of progeroid syndromes characterized by signs of aging and called HGPS-like. Most of HGPS-like patients carry mutations near the donor splice site of exon 11, causing the production of variable quantities of aberrant prelamin A isoforms. In particular, Prelamin A Δ90 transcript excludes the 270 nucleotides of exon 11 because of the abolition of the normal donor splice site. The resulting deletion is predicted to preserve the Prelamin A open reading frame (r.[=, 1699_1968del], p.(Gly567_Gln656del)). The mutation responsible for the production of Prelamin A Δ35 generates a conservative substitution of serine with threonine and activates a cryptic splice site resulting in the expression of a truncated prelamin A lacking 35 amino acids (r.[=, 1864_1968del], p.[Thr623Ser, Val622_Gln656del]) (Barthelemy et al., 2015; Harhouri et al., 2016). The group of progeroid syndromes includes also patients affected with Mandibuloacral Dysplasia type B (MAD-B), carrying a homozygous mutation in *ZMPSTE24*, encoding FACE1 protease involved in Prelamin A maturation and leading to accumulation of wild type farnesylated Prelamin A. We hypothesized that MG132 could also be beneficial for HGPS-like patients whose cells express WT prelamin A, Prelamin A Δ35, Prelamin A Δ50 or/and Prelamin A Δ90. Here, we test the effects of MG132 on HGPS-like and MAD-B cells and characterize the drug effect on aberrant prelamin A isoforms clearance as well as the improvement of cell phenotypes.

## RESULTS

### Patients’ Molecular and Clinical Features

Patients included in this study showed variable disease severity compared to classical HGPS but presented with similar phenotypes, including growth retardation, hair loss, prominent forehead, prominent superficial veins, thin skin, loss of subcutaneous fat and lipodystrophy (**Figure. 1A**). HGPS-like patients were previously reported to present distinct aberrant splicing patterns of Prelamin A pre-mRNAs due to mutations located around exon 11 donor splice site (**Figure. 1B, C**) (Harhouri et al., 2016). Briefly, patient HGPS-L1 carrying the *LMNA* heterozygous c.1968+2T>C mutation was referred to our center at the age of five years. She was diagnosed with the disease when she was 10 months old, presenting with a typical HGPS clinical phenotype, including frontal bossing, prominent veins on her scalp and forehead, sparse hair, micrognathism with delayed dentition, growth retardation (since birth, her length varied from the 2nd to the 10th centiles for age; the weight was stably <3rd centile for age), subcutaneous lipoatrophy, dry skin with pigmentary changes on the neck and trunk, acroosteolyses with onychodystrophy of hands and feet; laboratory findings have evidenced recurrent thrombocytosis (480–535 k/μL; normal values: 140–450 k/μL), elevated transaminases, glucose, calcium and phosphorus, as already observed in classical HGPS patients (Merideth et al., 2008). Patient HGPS-L2, carrying the heterozygous *LMNA* c.1968+1G>A mutation, showed a very similar progeroid laminopathy, though evolving more severely. She was diagnosed at nine months of age and already showed contractions of her ankles, knees, and wrist. She subsequently developed arthritis on several articulations. Her feeding was poor, and she had frequent constipation episodes. She had bilateral hip dislocation and at the age of three years she suffered from a femur fracture. At age six, she suffered from tachycardia together with sudden right arm paresis; cerebral CT scan/MRI showed multiple micro-infarcts, including recent and old ones, while echocardiography showed left ventricular thickening. After partial recovery from stroke, she suffered from a chest infection together with painful nails’ infections. Patients HGPS-L3 (*LMNA* heterozygous c.1968+5G>C), HGPS-L6 (*LMNA* heterozygous c.1868C>G) and HGPS-L5 (*LMNA* heterozygous c.1968G>A) were previously reported by Barthelemy et al. (Barthelemy et al., 2015). Patient HGPS-L4 (*LMNA* heterozygous c.1968+6C>T) is first reported in this work. She was referred to our clinics at age 4 years and presented with sparse hair and eyebrows, small chin, a thin nose, prominent nipples, dyspigmentation with hyper/hypo-pigmented areas, sclerodermatous changes on her chest. The MAD-B patient was first referred to us at age 6.5 years. She presented with growth retardation, exophtalmia, low-set ears, retro-micrognathism (mandibular hypoplasia), sparse hair and thin, dry skin with hypopigmented lesions, especially on the trunk. Subcutaneous lipoatrophy gave her a muscular pseudo-hypertrophic appearance. Molecular genetic diagnosis allowed the identification of a new homozygous mutation in the *ZMPSTE24/FACE1* gene’s exon 10: c.1274T>C, p.(Leu425Pro), confirming the B-type Mandibuloacral dysplasia phenotype in the patient (Ben Yaou et al., 2011; Harhouri et al., 2016).

**Figure. 1.**
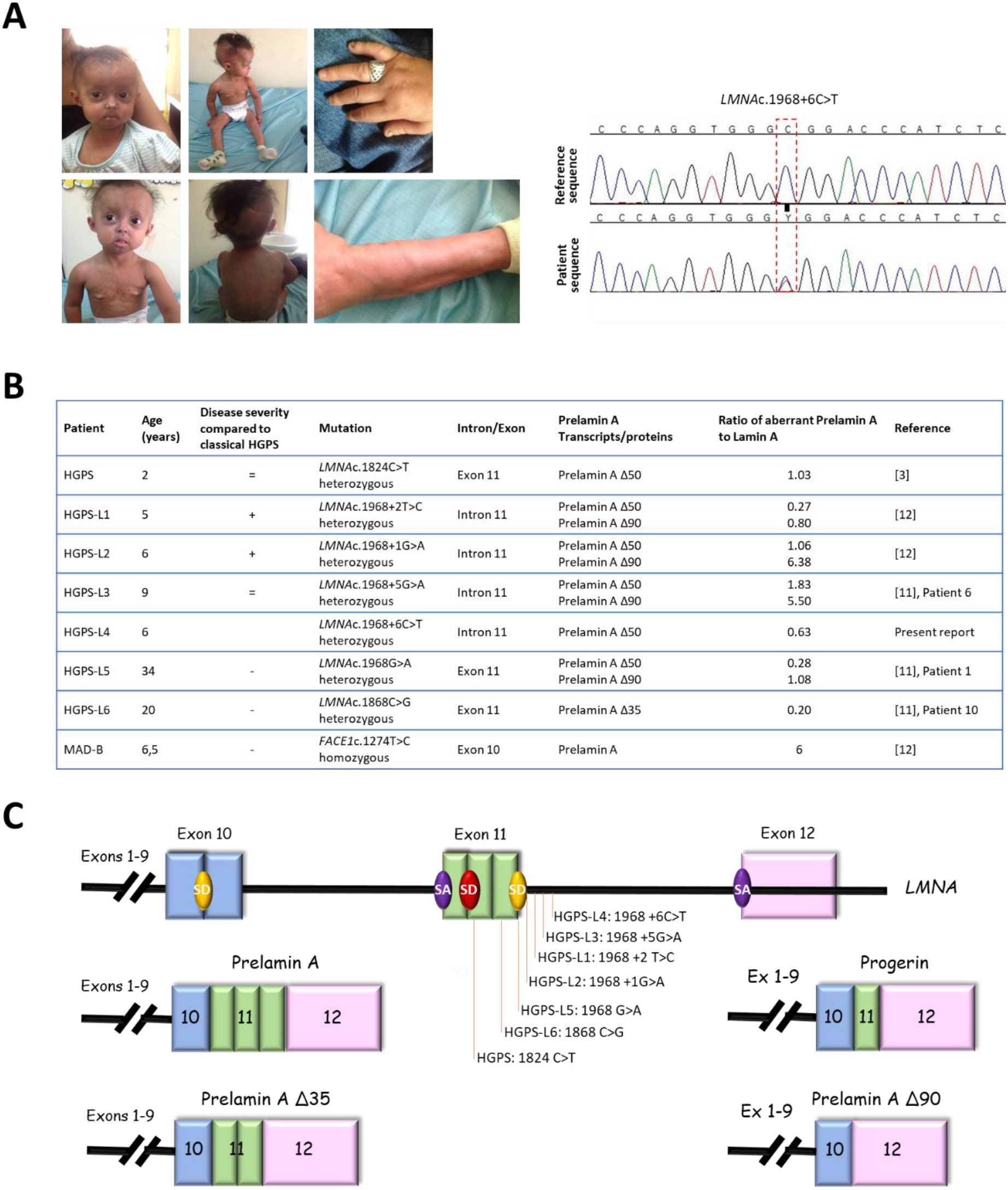
Clinical and molecular description of cell lines. (**A**) Pictures and electropherograms of patient HGPS-L4 at age 6 years showing progeroid features including sparse hair and eyebrows, small chin, a thin nose, prominent nipples, dyspigmentation with hyper/hypo-pigmented areas and sclerodermatous changes on her chest. The heterozygous c.1968+6C>T LMNA mutation was confirmed by Sanger sequencing (**B**) Characterization of LMNA and FACE1 (ZMPSTE24) gene mutations in HGPS-like and MAD-B patients eliciting aberrant Prelamin A splicing or wild type Prelamin A accumulation. Variable disease severities compared to classical HGPS are indicated with “+”: more, “−”: less, or “=”: equal severity. The ratios of aberrant Prelamin A to Lamin A isoforms are shown, issued from Western Blot data, except for Prelamin A against which no antibodies are available and so were determined based on the transcript levels. (**C**) Locations of LMNA mutations and schematic representation of the aberrant Prelamin A isoforms. SD: Splice Donor site, SA: Splice Acceptor site.

### MG132 reduces aberrant prelamin A levels in HGPS-like and MAD-B fibroblasts

Our previous studies in typical HGPS cells have shown that MG132 promotes both progerin degradation through autophagy activation and reduction of progerin synthesis mediated by regulation of SRSF-1 and SRSF-5, playing opposite role in the utilization of the *LMNA* and progerin 5’ splice site. Therefore, hypothesizing that MG132 might have the same effects on aberrant prelamin A isoforms clearance in HGPS-like cells, we performed quantitative reverse transcription–polymerase chain reaction (RT-PCR) assays using primers specific for prelamin A mRNA isoforms in MG132- and DMSO-treated HGPS-like cells. As shown in **Figure. 2A**, when compared to DMSO-treated cells, MG132 treatment at 500 nM for 24h induces aberrant prelamin A mRNA downregulation, suggesting that the drug acts at the transcriptional levels. Indeed, we observed significant reductions in prelamin A Δ50 and prelamin A Δ90 mRNAs in HGPS-L1, HGPS-L2, HGPS-L3 and HGPS-L5 patients’ cells, prelamin A Δ50 mRNA in HGPS-L4 and prelamin A Δ35 mRNA in HGPS-L6 patient cell. The treatment also significantly decreased the production of lamin A transcripts in MAD-B fibroblasts. To further evaluate the MG132-associated decrease in prelamin A isoforms at the protein levels, we treated HGPS-like fibroblasts with 500 nM MG132 for 48h. Quantification of the Western blotting experiments revealed clear reductions in prelamin A Δ50 in HGPS-L1, HGPS-L2, HGPS-L3, HGPS-4 and HGPS-L5 patients’ cells, and prelamin A Δ35 in HGPS-L6 patient cell. In MAD-B cells, the treatment also significantly decreased the production of prelamin A (**Figure. 2B**). Interestingly, in all the tested HGPS-like cell lines and concomitantly with the decrease of aberrant prelamin A levels, LC3B-I to LC3B-II autophagic switch was increased.

**Figure. 2.**
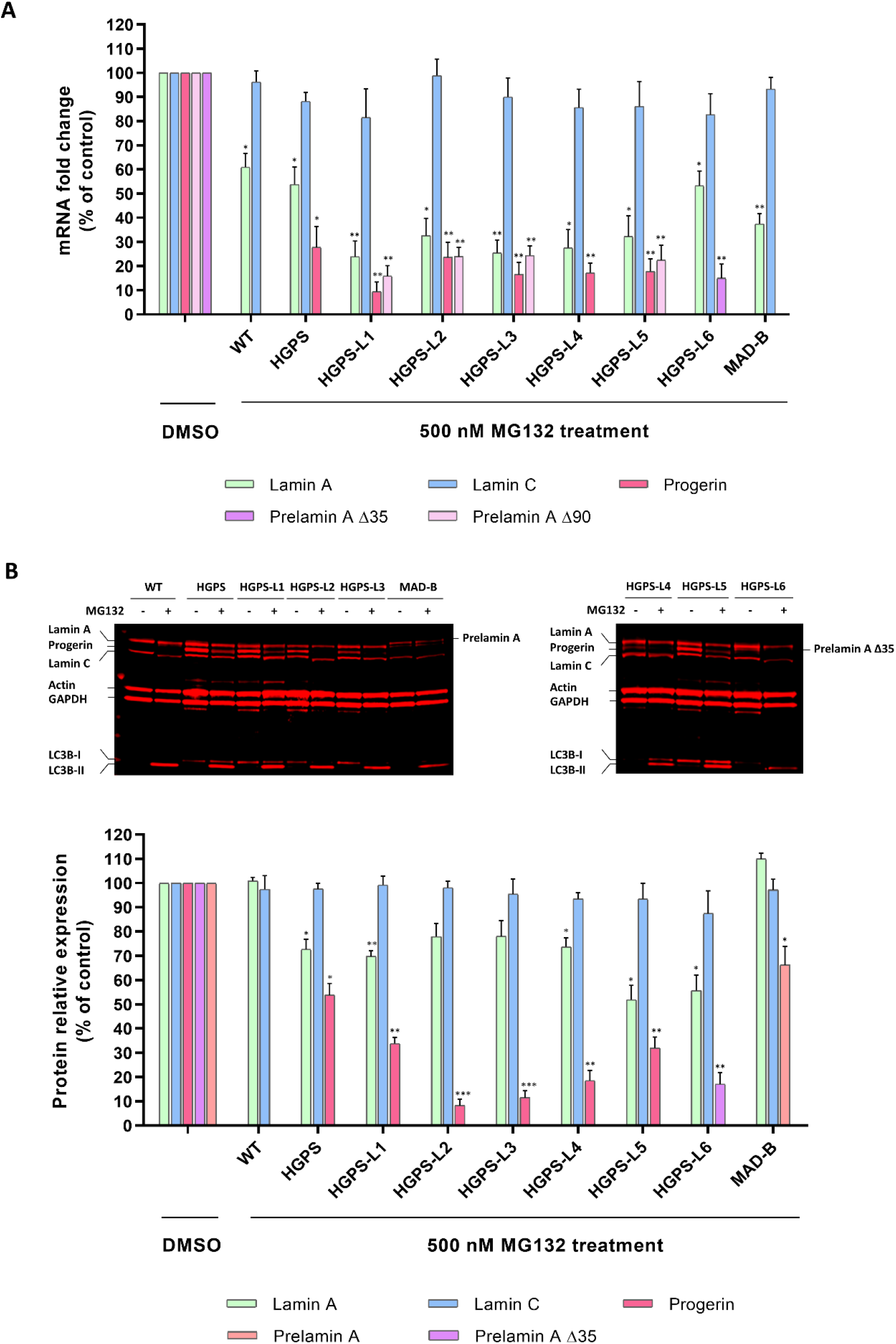
MG132 reduces aberrant prelamin A levels in HGPS-like and MAD-B fibroblasts. (**A**) Downregulation of aberrant prelamins A transcripts (Δ50: progerin, Δ35, Δ90 and WT prelamin A) in response to MG132. Quantitative real-time PCR analyses of lamin A, prelamin A Δ50 (progerin), Prelamin A Δ35, Prelamin A Δ90, Lamin C and RPS13 mRNA levels in HGPS, HGPS-like, MAD-B and WT fibroblasts treated for 24h with 500 nM MG132 relative to DMSO-treated cells (Control). The fold change of each transcript was determined by normalizing its value to that of RPS13 for each condition. (mean ± SEM, n = 4, Student’s t-test, *p < 0.05, **p < 0.01, experimental vs. control). (**B**) MG132 reduce aberrant Prelamin A protein levels in HGPS-like and MAB-B patient cells. Upper panels: Western blotting evaluation of Lamin A/C, Progerin, Prelamin A, Prelamin A □35 in whole lysates from WT, HGPS, HGPS-like and MAD-B fibroblasts treated with DMSO (-), 500 nM MG132 for 48h (+). Lower panels: Lamin A/C, Progerin, Prelamin A, Prelamin A □ 35 expression levels were normalized to GAPDH values using ImageJ software. (mean ± SEM, n = 3, Student’s t-test, * p < 0.05, ** p < 0.01, *** p < 0.001. MG132-treated vs. DMSO-treated cells).

### MG132 reduces senescence, enhances proliferation and migration in HGPS-like and MAD-B patient cells

In primary fibroblasts from HGPS patients, progerin accumulation results in premature senescence, a major hallmark of HGPS, as well as of normal aging cells (Goldman et al., 2004;Scaffidi & Misteli, 2005; Vidak & Foisner, 2016). Therefore, we hypothesized that MG132-induced clearance of progerin might delays senescence in HGPS-like cells. To test MG132 efficacy, we first measured senescence by quantification of a luminescent signal that is dependent on and correlates with β-galactosidase activity. Interestingly, all HGPS-like cells treated with 500 nM MG132 for 96 h exhibited a decreased senescence rate (**Figure. 3A**). Furthermore, using Senescence Associated β-galactosidase staining, we observed that this MG132 treatment scheme induces a decrease in the number of senescent cells when compared to DMSO-treated cells (**Figure 3B**).

**Figure.3.**
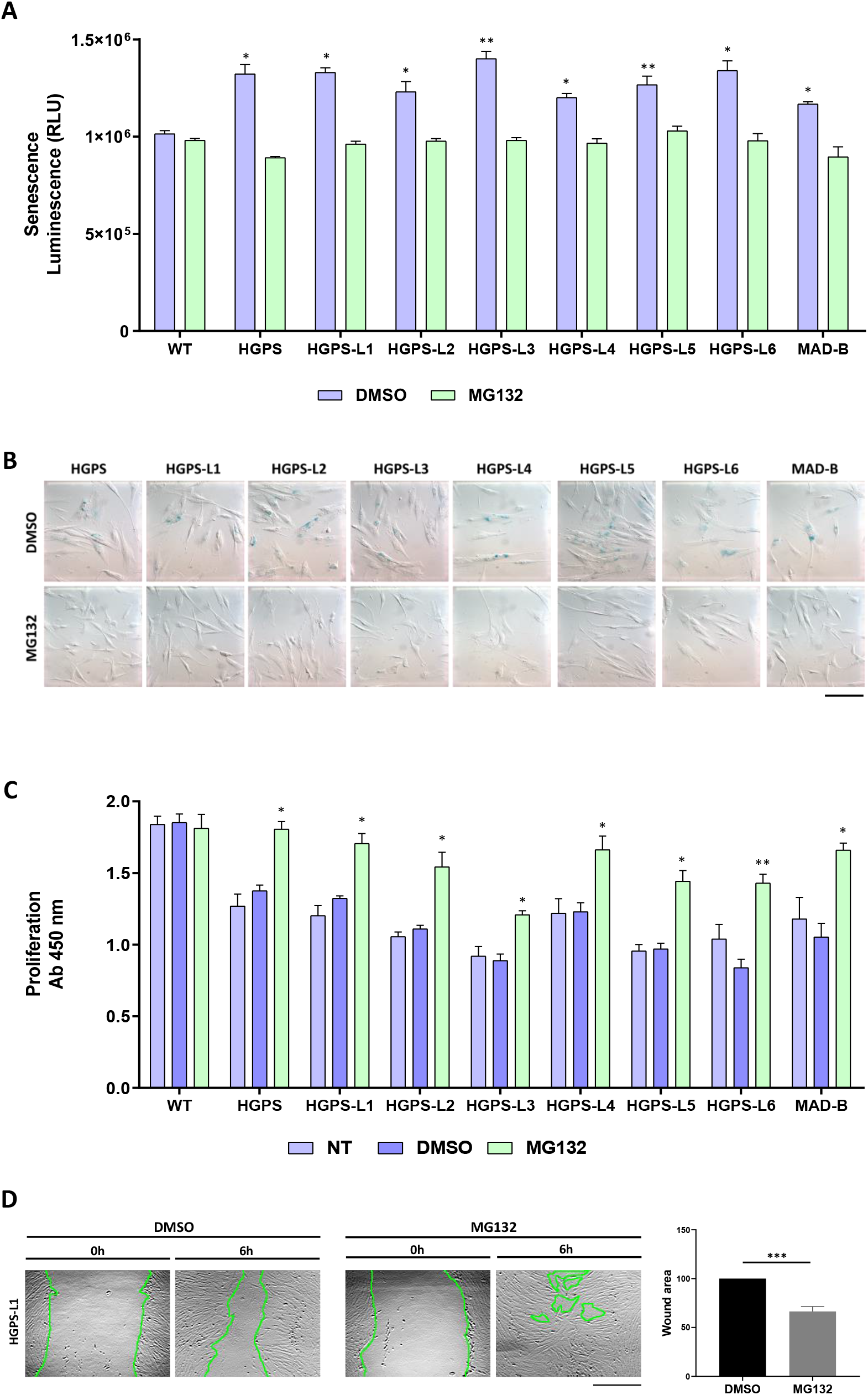
MG132 reduces senescence, enhances proliferation and migration in HGPS-like and MAD-B patients’ cells. (**A**) Luminescence-based quantification of senescence rate in WT, HGPS-like and MAD-B fibroblasts treated with 500 nM MG132 for 96 h relative to DMSO-treated cells. Each experiment was performed on cells at the same passage level. Senescence is determined as relative light units (RLU). (mean ± SEM, n = 3, Student’s t-test, *p < 0.05, ** p < 0.01. MG132-treated vs. DMSO-treated cells). (**B**) Colorimetric detection of senescence-associated β galactosidase in HGPS-like and MAD-B fibroblasts treated with 500 nM MG132 for 96 h relative to DMSO-treated cells. Each experiment was performed on cells at the same passage level. β-galactosidase blue staining is lower in cells treated with MG132 compared to cells treated with DMSO. (**C**) Cell proliferation rate based on the incorporation of bromodeoxyuridine (BrdU) into the DNA was expressed as absorbance OD 450 nm. (**D**) Left panel: an example of wound healing assay performed on HGPS-L1 fibroblasts treated for 6h with DMSO or MG132 (500 nM). Right panel: the results of wound healing assays on individual samples (Figure S1) were grouped into biological replicates (1 HGPS, 6 HGPS-like and 1 MAD-B) to perform statistical tests. (mean ± SEM, n = 8, Student’s t-test, *** p < 0.001. MG132-treated vs. DMSO-treated cells). Results are expressed as a percentage of the area of the original wound, and normalized to DMSO-treated cells, considered as 100%. Scale bar, 100 μm.

Primary fibroblasts from HGPS patients exhibit proliferative defects (Goldman et al., 2004). To determine whether MG132-induced clearance of progerin has any beneficial effects on cell proliferation, we examined the proliferation rates of HGPS-like fibroblasts with MG132 or DMSO treatment and found that in all the tested cell lines, proliferation rates were increased by a 96h MG132 treatment at 500 nM, when compared to the DMSO-treated cells (**Figure. 3C**). As in HGPS (Chang et al., 2019), nuclear architecture and cell migration are impaired during physiological aging (Pienta & Coffey, 1990). We investigated the effect of MG132 treatment on cell migration. “Wound-healing” assays showed that most of the MG132-treated HGPS-like and MAD-B cells (6/8) were able to migrate and to “heal the wounds” better than their control DMSO-treated counterparts (**Figure 3D and Figure S1**).

### MG132 treatment rescues the level of proteins whose expression is altered in HGPS-like and MAD-B cells

Other characteristics of fibroblasts from individuals with HGPS cells include a loss of peripheral heterochromatin and downregulated tri-methyl lysine 9 of core histone H3 (Tri-Me-K9) (Scaffidi & Misteli, 2005; Vidak & Foisner, 2016) as well as reduced levels of the nuclear components, lamin B1, and lamina-associated polypeptide (LAP2α) (Goldman et al., 2004;Scaffidi & Misteli, 2005). By immunocytochemistry studies, as observed in (**Figure. 4**), levels of these proteins are also reduced in HGPS-like and MAD-B cells compared to WT, supporting a negative correlation between aberrant prelamin A accumulation and down-regulation of several nuclear proteins, including histone modification patterns. Importantly, treatment with MG132 restored the levels of histone H3-Tri-Me-K9, lamin B1 and LAP2α in most cells and normalizes their expression to levels close to those of normal fibroblasts.

**Figure. 4.**
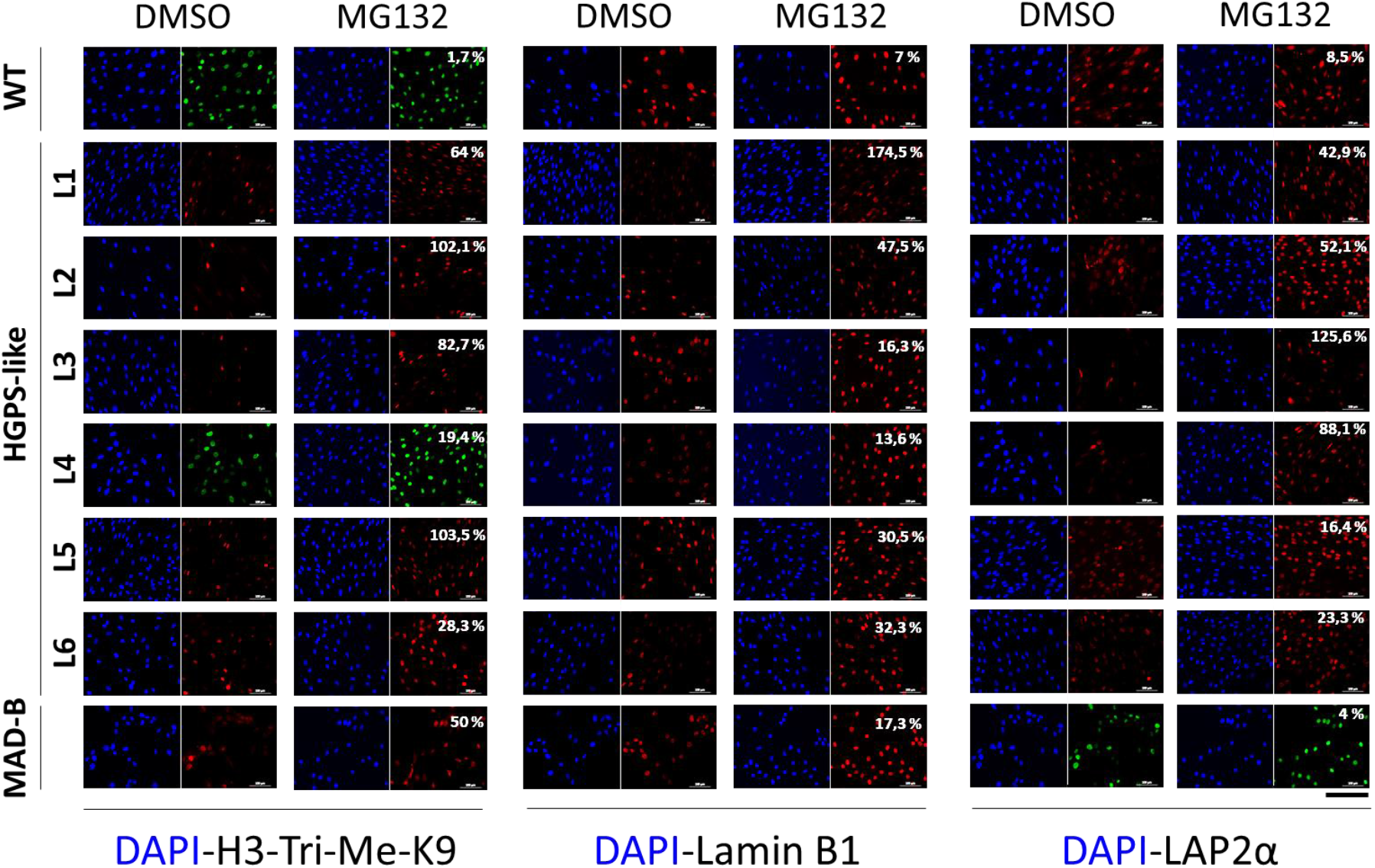
MG132 treatment rescues the level of proteins whose expression is altered in HGPS-like and MAD-B cells. Immunofluorescence microscopy on primary dermal fibroblasts from a healthy individual (WT), HGPS-like and MAD-B patients, treated with 500 nM MG132 or equal volume of DMSO for 48h. Cells were stained with DAPI (blue) and antibodies to tri-methyl lysine 9 of core histone H3 (H3-Tri-Me-k9), Lamin B1 and LAP2α. The percentage of positive staining is indicated, the percentage indicates the variations in signal intensity between MG132-treated cells and control, each normalized to the corresponding nuclei number, at least 200 fibroblast nuclei were randomly selected for each cell line (n = 3) and examined using ImageJ software. Scale bar, 200 μm.

### Treatment of HGPS-like and MAD-B cells with MG132 reduces the levels of DNA damage

Previous studies have shown that HGPS cells accumulate defective DNA damage response (DDR) playing a key role in the premature aging phenotypes (Gonzalo & Kreienkamp, 2015;Scaffidi & Misteli, 2006). Progerin causes chromatin perturbations, specially, the global loss of histone H3-Tri-Me-K9, leading to the formation of DSBs (double-strand breaks) and abnormal DDR as evidenced by the accumulation of phosphorylated histone γ-H2AX foci and impaired recruitment of p53-binding protein 1 (53BP1) to sites of DNA damage (Liu et al., 2005; Zhang et al., 2016). We performed γ-H2AX/53BP1 double immunofluorescence staining and observed more γ-H2AX-positive foci in HGPS-like and MAD-B cells than those observed in control, with defective recruitment of 53BP1 to these sites (**Figure. 5**). However, MG132 treatment reduced the number of nuclei with γ-H2AX foci. Moreover, we observed more effective recruitment of 53BP1 to the remaining γ-H2AX foci.

**Figure. 5.**
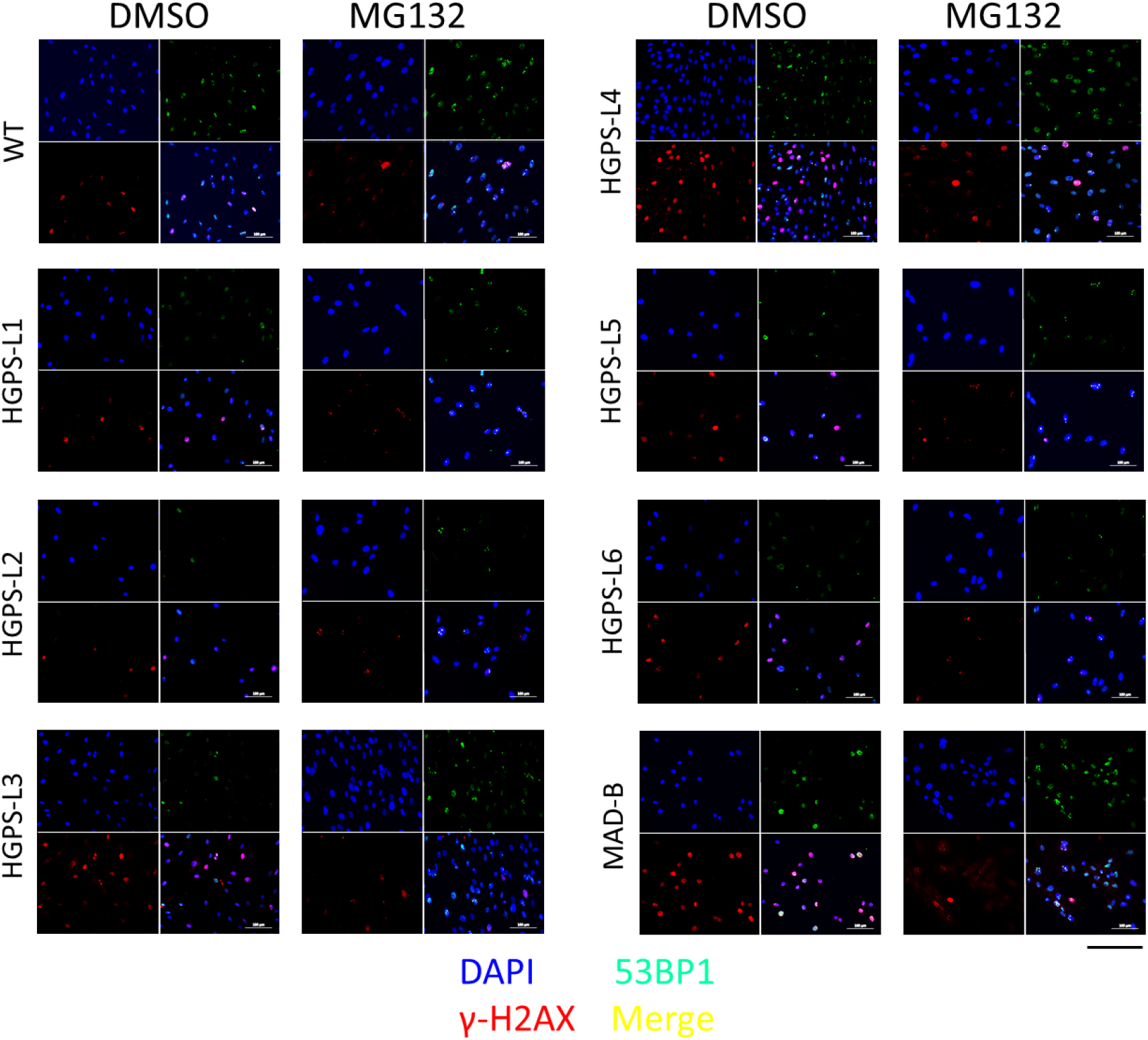
Treatment of HGPS-like and MAD-B cells with MG132 reduces the levels of DNA damage. Immunofluorescence microscopy on primary dermal fibroblasts from a healthy individual (WT), HGPS-like and MAD-B patients, treated with 500 nM MG132 or equal volume of DMSO for 48h. Cells were stained with DAPI (blue) and antibodies to the indicated proteins. Scale bar, 200 μm.

### Anti-inflammatory effects of MG132 in HGPS-like and MAD-B cells

Many altered signaling pathways have been described in HGPS cells (Harhouri et al., 2018;Vidak & Foisner, 2016). Among them, hyperactivation of NF-κB inflammatory pathway (Osorio et al., 2012). In a previous study, crossing a mouse model for premature aging, *Zmpste*24^-/-^, with transgenic mice displaying reduced NF-κB signaling, extends longevity and prevents the development of progeroid features. Moreover, inhibition of NF-κB by sodium salicylate efficiently prevents the disease phenotypes in *Zmpste*24-deficient mice and extends longevity in the HGPS model, *Lmna*^G609G/G609G^(Osorio et al., 2012). On the other hand, MG132 is also known to attenuate the degradation of NF-κB inhibitor, I-κB **(Figure S2)**, resulting in the inhibition of proinflammatory cytokines secretion (Hozhabri, Kim, & Varanasi, 2014;Mathes, O’Dea, Hoffmann, & Ghosh, 2008; Ortiz-Lazareno et al., 2008).

To further investigate the cellular inflammatory response of MG132-treated HGPS fibroblasts, and given that this cell type (fibroblasts from skin biopsy) is known to secrete high levels of inflammatory cytokines (Mahmoudi et al., 2019), we performed RNA-seq experiments (accession number: E-MTAB-5807) and analyzed the expression levels of NF-κB gene signatures in classical HGPS fibroblasts treated with MG132. Interestingly, we found a decrease in the transcript’s levels of TNFα, IL-6, IL-18, IL-19, IL-34, IL-1 receptor accessory, IFNα-R2, interferon regulatory factor 7, TGFβ-R3 and EGF-R, as well as the increase of other anti-inflammatory transcripts: IL-1R2, IL-1R antagonist, NF-κB inhibitor α, NFκB inhibitor β, NFκB repressing factor and NF-κB inhibitor like 1 **(Figure S3)**. In order to assess the inflammatory response with MG132 on HGPS-like and MAD-B fibroblasts, we performed quantitative real-time PCR using selected inflammatory genes expression arrays on culture supernatants of fibroblasts treated with MG132, TNFα alone and in combination. As described in **Figure. 6A and Figure S4**, we found that MG132 reduces the transcript levels of proinflammatory cytokines (IL-1α, IL-1β, IL-6, TNFα) in HGPS-like and MAD-B patient cells. Moreover, treatment with MG132 reduced the transcript levels of proinflammatory mediators induced by recombinant TNFα (IL-1α, IL1-β, IL-6, IL-8, TNFα, IFNβ1, EGF-R, NFκB1, NFκB2, RelA). In the same way, using ELISA, we found a significative downregulation of several proinflammatory cytokine, such as IL-1β, Il-6, IL-17A, TNFα, TGFβ and CXCL1. Again, MG132 reduces TNFα-induced secretion of the proinflammatory cytokines IL-1β, Il-6, TNFα, IFNγ and TGFβ (**Figure. 6B and Figure S5**).

**Figure. 6.**
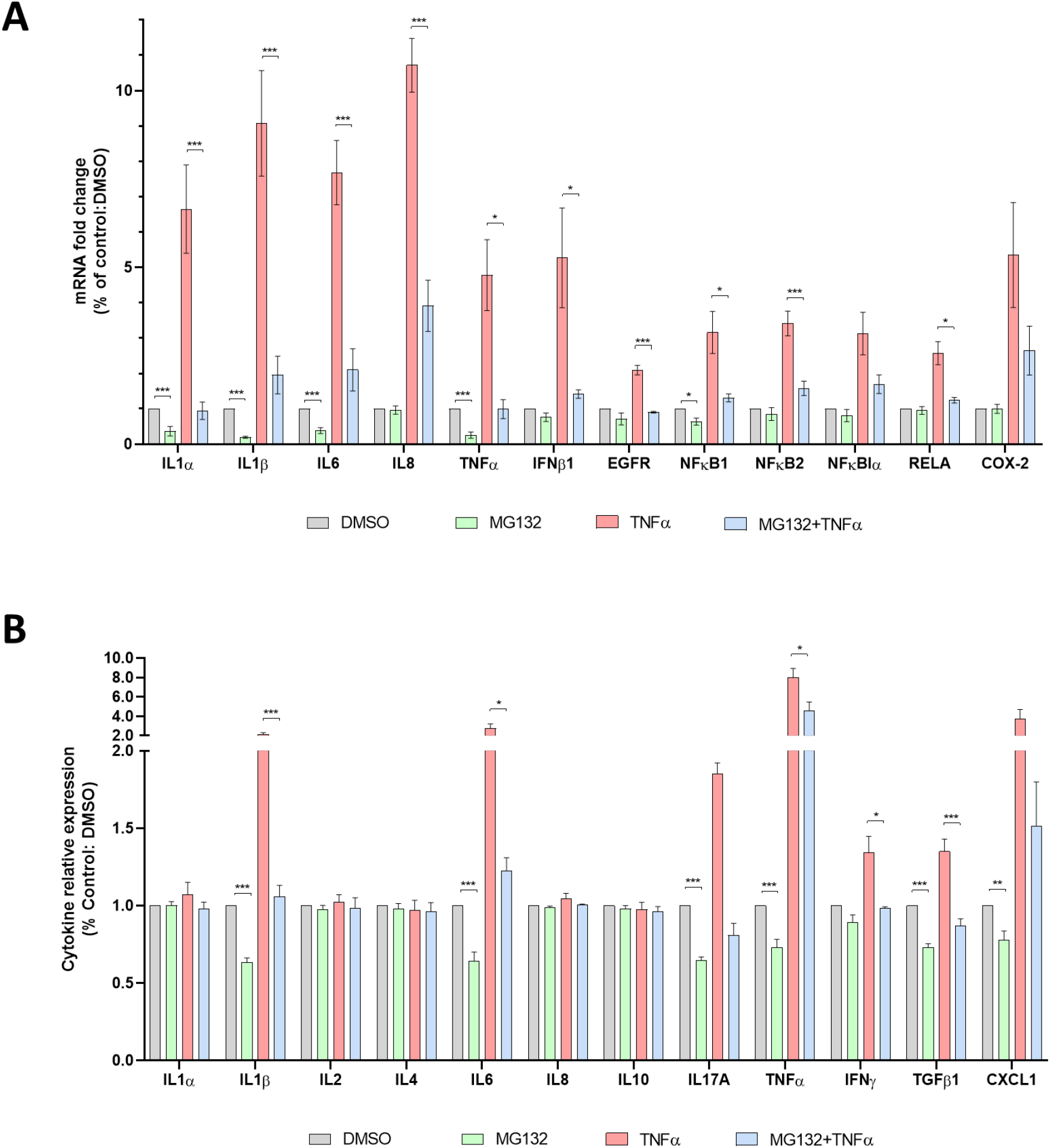
Anti-inflammatory effects of MG132 in HGPS-like and MAD-B cells. (**A**) Quantitative real-time PCR using selected inflammatory genes expression arrays in culture supernatants of HGPS, HGPS-like and MAD-B fibroblasts treated for 6h with MG132 (500 nM), TNFα (10 ng/ml) alone and in combination or DMSO as a vehicle control. The results of individual samples (Figure S3) were grouped into biological replicates (1 HGPS, 6 HGPS-like and 1 MAD-B) to perform statistical tests. (mean ± SEM, n = 8, Student’s t-test, * p < 0.05, **p < 0.01, ***p < 0.001. MG132-treated vs. DMSO-treated cells and MG132 + TNFα-treated vs. MG132-treated cells). (**B**) Enzyme-Linked Immunosorbent Assay (ELISA) using multi-analyte ELISA arrays to measure inflammatory cytokines in culture supernatants from HGPS, HGPS-like and MAD-B fibroblasts treated for 24h with MG132 (500 nM), TNFα (10 ng/ml) alone and in combination or DMSO as a vehicle control. The results of individual samples (Figure S4) were grouped into biological replicates (1 HGPS, 6 HGPS-like and 1 MAD-B) to perform statistical tests. (mean ± SEM, n = 8, Student’s t-test, *p < 0.05, ** p < 0.01, *** p < 0.001. MG132-treated vs. DMSO-treated cells and MG132+TNFα-treated vs. MG132-treated cells).

## DISCUSSION

We previously showed that the benefit of MG132 on classical HGPS fibroblasts and mice is mediated by 1/ induced macroautophagy leading to progerin degradation and 2/ blocking progerin production by reducing SRSF-1 expression levels and increasing expression levels of SRSF-5, controlling aberrant splicing of prelamin A precursor mRNA. MG132-induced clearance of progerin is partly due to autophagy activation in HGPS cells that is supported by the increased amounts of LC3B-II/LC3B-I ratios on immunoblotting assays, progerin delocalization into cytoplasmic autophagic vacuoles, increased autophagic transcript levels using RNA-seq experiments and partial restoration of progerin levels in presence of chloroquine or bafilomycin A1 which are commonly used as autophagy inhibitors. Otherwise, it is known that impairment of the Ubiquitin-proteasome system is compensated by activation of autophagy (X. J. Wang et al., 2013; Zang et al., 2012; Zhu, Dunner, & McConkey, 2010). MG132 treatment improves HGPS fibroblast phenotypes, reduces cell senescence, and improves their viability and proliferation. Injection of MG132 into the skeletal muscle of our progeria mice model (*Lmna*^G609G / G609G^) locally reduces progerin and SRSF-1 expression levels (Harhouri et al., 2017).

*LMNA* mutations other than the classical c.1824C>T (p.G608G) have been shown to cause the production of progerin and/or other truncated or wild type prelamin A isoforms in patients affected with HGPS-like and MAD-B syndromes (Barthelemy et al., 2015; Moulson et al.,2007). We therefore hypothesized that MG132 could also have a beneficial impact on those cells, since they share the same pathophysiological mechanism based on abnormal splicing of prelamin A pre-mRNA. To this end, we evaluated the treatment’s efficacy of MG132 in reducing the production of all prelamin A isoforms, including aberrantly accumulated prelamin A either truncated (HGPS-like) or wild type (MAD-B). Here, we show a significant decrease of each aberrant transcript’s production (Prelamin A Δ35, Δ50 and Δ90) as well as the corresponding abnormal proteins. Interestingly, MG132 not only induces the synthesis blockade of the aberrant prelamin A isoforms and the clearance of the corresponding proteins already expressed, but also results in the improvement of several biological parameters including cellular senescence, proliferation, altered protein expression, DNA damage and repair as well as inflammatory cytokine expression.

The results of the present study demonstrate that MG132 is at least as efficient as morpholinos treatment (Harhouri et al., 2016) with a wide range of beneficial effects on HGPS and HGPS-like, potentially due to both an impressive clearance of aberrant prelamin A isoforms and rescue of downstream noxious cascades. Notably, MG132 treatment lowers the levels of mediators of the inflammatory pathways. In agreement with our study, MG132 is known to inhibit the secretion of proinflammatory cytokines and decrease of I-κB degradation, resulting in the abolition of NF-kB activation (Ahmed et al., 2017; Hozhabri et al., 2014; Ortiz-Lazareno et al.,2008). Inflammation is a major regulator of the physiological and premature aging process (Neves & Sousa-Victor, 2020). Moreover, the major clinical hallmark of progeria is atherosclerosis, leading to premature death by myocardial infarction or stroke (Capell, Collins, & Nabel, 2007; Hennekam, 2006; Stehbens, Delahunt, Shozawa, & Gilbert-Barness, 2001). These findings, together with the fact that arterial lesions in both typical atherosclerosis and HGPS exhibit inflammation, calcification and the loss of vascular smooth muscle cells (VSMCs) (Hamczyk et al., 2018; Olive et al., 2010) support the need of targeting inflammation signaling cascade for the treatment of premature aging disorders.

On the other hand, NRF2 antioxidant pathway has been described as a driver mechanism in HGPS, due to impaired NRF2 transcriptional activity and consequently increased chronic oxidative stress (Kubben et al., 2016). Importantly, it has been shown that MG132 activates NRF2-ARE signaling pathway, which is associated with increased Nrf2 transcription and expression, leading to prevention of oxidative stress induced both in cardiovascular and renal injury (Cui, Bai, Luo, Miao, & Cai, 2013) and in several human endothelial and vascular smooth muscle cells (Dreger et al., 2010; Miao et al., 2013). Furthermore, MG132 was reported to have a significant preventive and therapeutic effect on accelerated atherosclerosis in rabbits (Feng et al., 2010), diabetic cardiomyopathy in diabetic mouse model (Y. H. Wang et al., 2013), arthritis associated with joint inflammation in rats (Ahmed et al., 2010). Interestingly, these features are exhibited by HGPS patients who might benefit from the same treatment.

In order to evaluate the effects of MG132 in our Knock-in progeria mouse model (*Lmna*^G609G/G609G^), carrying the c.1827C>T (p.Gly609Gly) mutation, we previously showed that the reduction of progerin levels upon IV or IP systemic treatment was not significant, suggesting that the molecule is unstable when injected systemically. Therefore, we performed intramuscular injections in *Lmna*^G609G/G609G^ tibialis anterior muscle. In this case, treatment with MG132 induced a significant decrease of progerin and SRSF-1 levels, in the treated muscle as compared to the untreated contralateral muscle (Harhouri et al., 2017).

MG132 rapid catabolism upon IV or IP administration is a clear limiting step for systemic delivery of the drug. We therefore set up a collaboration with an academic laboratory to develop MG132-derivatives in order to optimize the chemistry of the molecule, to improve its stability and efficacy and to analyze and minimize adverse effects, aiming to obtain *in vivo* systemic efficacy on reversion of premature aging phenotypes in *Lmna*^G609G/G609G^ mice.

Altogether, the originality and therapeutic potential of MG132 for HGPS and related diseases is based on its triple mechanism of action: targeting progerin production and degradation, in combination with decreased downstream noxious effects. Here, we have provided evidence that the use of MG132 could be extended to other syndromes characterized by the accumulation of truncated or wild type prelamin A. Our results establish a preclinical proof of principle for the use of MG132 or its druggable derivatives in HGPS-like and MAD-B syndromes with a strong potential for clinical administration in future trials.

## MATERIALS AND METHODS

### Patients and Samples

Samples were collected from eight patients affected with typical HGPS, HGPS-like or MAD-B syndromes, showing different genomic pathogenic variants, variable clinical phenotypes and disease severity but all having in common the accumulation of aberrant and toxic Prelamin A isoforms. Patients were from USA (HGPS-L1), UK (HGPS-L2 and HGPS-L6), Greece (HGPS-L3), Nicaragua (HGPS-L4), France (HGPS-L5) and Togo (MAD-B). Informed consents were obtained from the patients or the parents of minor patients included in this work, allowing studies on their cells as part of a diagnosis and research program, complying with the ethical guidelines of the institutions involved. Parents also gave written consent for picture publication, including uncovered faces. The dermal fibroblast cell line from patient HGPS-L 1 was provided by the Progeria Research Foundation Cell and Tissue Bank under the cell line name PSADFN386 (www.progeriaresearch.org); the other human dermal fibroblast cell lines were issued from a skin biopsy, prepared and stored by the certified Biological Resource Center (CRB AP-HM Biobank; NF S96-900 & ISO 9001 v2015 Certifications), Department of Medical Genetics, La Timone Hospital of Marseille, according to the French regulation. The fibroblast cell lines used belong to a biological sample collection declared to the French Ministry of Health (declaration number DC-2008-429) whose use for research purposes was authorized by the French Ministry of Education, Research and Innovation (authorization number AC-2011-1312; AC-2017-2986).

### Genomic characterization of *LMNA* variants

All the patients included in this work, excepted patient HGPS-L4, were already described (Barthelemy et al., 2015; Harhouri et al., 2016). Patient HGPS-L4 is first described in this work and her genomic characterization was performed as described in (Barthelemy et al., 2015), upon Sanger sequencing of the *LMNA* gene, which was directly performed in a diagnosis setting upon clinical suspicion. Briefly, Primer-3 designed specific primers were used for PCR amplification of each *LMNA* coding exon. PCR products were examined by agarose gel electrophoresis and then subjected to Sanger sequencing. Sequencher 4.8 (Gene Codes Corp., Ann Arbor, USA) was used for the interpretation of sequence variants. Sequence variants are described following the Human Genome Variations Society Guidelines available at https://varnomen.hgvs.org/. *LMNA* and *ZMPSTE24/FACE1* variants are respectively described relative to transcript reference sequences NM_170707.3 and NM_005857.5.

### Cell culture

Human dermal fibroblasts (established from a skin biopsy) were cultured in Dulbecco’s modified Eagle’s medium (Life Technologies) supplemented with 15% fetal bovine serum (Life Technologies), 2 mM L-glutamine (Life Technologies), and penicillin–streptomycin (Life Technologies) at 37°C in a humidified atmosphere containing 5% CO2. Testing for mycoplasma contamination was performed monthly. Fibroblasts were treated with media containing 500 nM or 5μM MG132 (474790, Merck Chemical LTD), 10ng/ml TNFα (210-TA, R&D systems), combination of 500 nM MG132 and 10ng/ml TNFα or with media containing the same volume of DMSO (vehicle control). The experiments were performed on fibroblasts of patients and healthy subjects matched for age and passage number.

### RNA sequencing (ArrayExpress accession number: E-MTAB-5807)

RNA sequencing was performed by IntegraGen (Evry, France). RNA samples were used to generate sequencing libraries with the TruSeq Stranded mRNA Sample Prep’ Illumina^®^. The libraries were sequenced on an Illumina HiSeq 4000 sequencer, yielding approximately 35 million 2 × 75-bp paired-end reads.

#### Quality control

Quality of reads was assessed for each sample using FastQC (http://www.bioinformatics.babraham.ac.uk/projects/fastqc/).

#### Sequence alignment and quantification of gene expression

A subset of 500,000 reads from each Fastq file was aligned to the reference human genome hg38 with TopHat2 to determine insert sizes with Picard. Full Fastq files were aligned to the reference human genome hg38 with TopHat2 (-p 24 -r 150 -g 2 -library-type fr-firststrand). Reads mapping to multiple locations removed. Gene expression was quantified using two non-overlapping transcriptome annotations: the full Gencode v25 annotation as well as a complementary lncRNA annotation. HTSeq was used to obtain the number of reads associated to each gene in the Gencode v25 database (restricted to protein-coding genes, antisense and lincRNAs) and to each gene in the additional lncRNA database. The Bioconductor DESeq package was used to import raw HTSeq counts for each sample into R statistical software and extract the count matrix. After normalizing for library size, the count matrix was normalized by the coding length of genes to compute FPKM scores (number of fragments per kilobase of exon model and millions of mapped reads). Bigwig visualization files were generated using the bam2wig python script. Unsupervised analysis: The Bioconductor DESeq package was used to import raw HTSeq counts into R statistical software, to obtain size factors, and to calculate a variance stabilizing transformation (VST) from the fitted dispersion–mean relations to normalize the count data. The normalized expression matrix from the 1,000 most variant genes (based on standard deviation) was used to classify the samples according to their gene expression patterns using principal component analysis (PCA) and hierarchical clustering.

#### Differential expression analysis

The Bioconductor DESeq package was used to import raw HTSeq counts into R statistical software, to obtain size factors and dispersion estimates and to test differential expression. Only genes expressed in at least one sample (FPKM ≥ 0.1) were tested to improve the statistical power of the analysis. A q-value threshold of ≤ 0.05 was applied to define differentially expressed genes.

### RNA isolation, reverse transcription, and real-time PCR

Total RNA was isolated using the RNeasy plus extraction kit (Qiagen, Valencia, CA, USA) and the samples were quantified and evaluated for purity (260 nm/280 nm ratio) with a NanoDrop ND-1000 spectrophotometer. 1 μg of RNA was reverse transcribed using SuperScript IV Reverse Transcriptase Kit (Thermo Fisher Scientific, Waltham, MA, USA). Real-time PCR amplification was carried out with the TaqMan Gene Expression Master Mix (Thermo Fisher Scientific, Waltham, MA, USA) on a LightCycler 480 (Roche, Germany) using predesigned primers for RPS13 (hs-01011487_g1), progerin (F: ACTGCAGCAGCTCGGGG. R: TCTGGGGGCTCTGGGC and probe: CGCTGAGTACAACCT), lamin A (F: TCTTCTGCCTCCAGTGTCACG. R: AGTTCTGGGGGCTCTGGGT and probe: ACTCGCAGCTACCG), and lamin C (F: CAACTCCACTGGGGAAGAAGTG. R: CGGCGGCTACCACTCAC and probe: ATGCGCAAGCTGGTG), Prelamin A Δ90 (F: CGAGGATGAGGATGGAGATGA. R: CAGGTCCCAGATTACATGATGCT, overlapping exons 10 and 12 and probe: CACCACAGCCCCCAGA) and Prelamin A Δ35 (F: ACTGCAGCAGCTCGGGG. R: AGTTCTGGGGGCTCGTGAC Probe: CGCTGAGTACAACCT) (Applied Biosystems). The gene expression of IL-1α, IL-1β, IL-6, IL-8, TNFα, IFN-β, EGFR, NFκB1, NFκB2, NFκBIα, Rel A, Cox-2 and the 18S rRNA control was assessed through real-time PCR using TaqMan^®^ Gene Expression Array Plates (ThermoFisher Scientific) containing predesigned, gene-specific primers and probes (Table 1). All qPCRs were performed using the program: UNG incubation at 50°C for 2 min, initial denaturation at 95°C for 10 min, 40 cycles of amplification: denaturation at 95°C for 15 s and annealing at 60°C for 1 min. All PCRs were performed in triplicate. Threshold cycle (Ct) values were used to calculate relative mRNA expression by the 2-ΔΔCt relative quantification method with normalization to RPS13 expression.

**Table 1.** List of genes used in real-time PCR using inventoried TaqMan Gene Expression Arrays.

| Gene | ID |
| --- | --- |
| 18s rRNA | Hs99999901_s1 |
| IL-1α | Hs00174092_m1 |
| IL-1β | Hs01555410_m1 |
| IL-6 | Hs00174131_m1 |
| IL-8 | Hs00174103_m1 |
| TNFα | Hs00174128_m1 |
| IFN-β1 | Hs01077958_s1 |
| EGFR | Hs01076090_m1 |
| NFκB1 | Hs00765730_m1 |
| NFκB2 | Hs01028890_g1 |
| NFκBIα | Hs00355671_g1 |
| RELA | Hs01042014_m1 |
| COX-2 | Hs00153133_m1 |

### Western blot

Total fibroblast proteins were extracted in 200 μl of NP40 Cell Lysis Buffer (Thermo Fisher Scientific, Waltham, MA, USA) containing Protease and Phosphatase Inhibitor Cocktail (Thermo Fisher Scientific, Waltham, MA, USA). Cells were sonicated twice (30s each), incubated at 4°C for 30 min and then centrifuged at 10,000 g for 10 min. Protein concentration was evaluated with the bicinchoninic acid technique (Pierce BCA Protein Assay Kit), absorbance at 562 nm is measured using nanodrop 1000 (Thermo Fisher Scientific). Equal amounts of proteins (40 μg) were loaded onto 10% Tris-glycine gel (CriterionTM XT precast gel) using XT Tricine Running Buffer (Bio-Rad, USA). After electrophoresis, gels were electro transferred onto Immobilon-FL polyvinylidene fluoride membranes (Millipore), blocked in odyssey Blocking Buffer diluted 1:1 in PBS for 1 h at room temperature, and incubated overnight at 4°C or 2 h at room temperature with various primary antibodies. Blots were washed with TBS-T buffer [20 mM tris (pH 7.4), 150 mM NaCl, and 0.05% Tween 20] and incubated with 1:10,000 IR-Dye 800-conjugated secondary donkey anti-goat or IR-Dye 700-conjugated secondary anti-mouse antibodies (LI-COR Biosciences) in odyssey blocking buffer (LI-COR Biosciences). For IR-Dye 800 and IR-Dye 700 detection, an odyssey Infrared Imaging System (LI-COR Biosciences) was used. GAPDH or actin were used as a total cellular protein loading control.

### Fluorescence microscopy

Fibroblasts were seeded into 4-well cell culture slides (Lab-tek, SPL Life Sciences), fixed with 4% paraformaldehyde, washed with PBS and permeabilized with 0.5% Triton X-100 for 15 min. After PBS washing, slides were incubated with 1% bovine serum albumin for 30 min at room temperature before adding the primary antibodies for 3 h at 37°C or overnight at 4°C. After washing, the cells were then incubated with secondary antibodies (A11001, A11058, Life Technologies; 1/400) for 1 hour at room temperature. Nuclei were stained with DAPI (50 ng/ml) diluted in Vectashield (Abcys) for 10 min at RT. The stained cells were observed with a Zeiss LSM 800 Confocal Microscope using Zen 2.3 software. All antibodies were tested in individual staining reactions for their specificity. Controls without primary antibody were all negative.

### Antibodies

Antibodies used in the study included: a rabbit anti-lamin A/C polyclonal antibody which reacts with lamin A, lamin C, and progerin (#SC-20681, used at 1:1,000 dilution for the Western blot analyses, Santa Cruz Biotechnology Inc.); a goat anti-prelamin A polyclonal antibody (#sc-6214 used at 1:1,000 dilution for the Western blot analyses, Santa Cruz Biotechnology Inc.); a mouse anti-actin monoclonal antibody (#MAB1501R, used at 1:5,000 dilution for the Western blot analyses, Merck Inc.); a mouse anti-glyceraldehyde-3-phosphate dehydrogenase monoclonal antibody (#MAB374, used at 1:10,000 for the Western blot analyses, Merck Millipore); a rabbit anti-LC3B polyclonal antibody (#2775, used at 1:1,000 for the Western blot analyses, Cell Signaling Technology); a rabbit anti-histone H3 (Tri-Me-K9) polyclonal antibody (#ab8898, used at 1:100 for immunofluorescence labeling, Abcam); a rabbit anti-lamin-B1 polyclonal antibody (#ab 16048, used at 1:100 for immunofluorescence labeling, Abcam); a rabbit anti-LAP2a polyclonal antibody (#ab5162, used at 1:100 for immunofluorescence labeling, Abcam); a mouse anti-γH2A.X (phospho S139) monoclonal antibody (#ab26350, used at 1:200 for immunofluorescence labeling, Abcam); a rabbit anti-53BP1 polyclonal antibody (#NB100-304, used at 1:1000 for immunofluorescence labeling, Novus Biologicals).

### Measurement of senescence

Senescence was measured using 2 assays: 1/ Beta-Glo Assay kit (Promega # E4720), according to the manufacturer’s instructions and utilizing a luciferin-galactoside substrate (6-O-β galactopyranosylluciferin). This substrate is cleaved by β-galactosidase to form luciferin and galactose. The luciferin is then utilized in a firefly luciferase reaction to generate a bright luminescent signal determined as RLUs using a GloMax-Multi Detection System: Luminometer (Promega, USA). 2/ Colorimetric detection of senescence-associated β galactosidase following the manufacturer’s protocol (Cell Signaling #9860). Cells were seeded in 4 chamber-wells slides (SPL Lifesciences, Korea), washed with PBS and fixed in Fixative solution (1/10 dilution) for 15 min at RT. Cells were washed in PBS, and stained overnight at 37°C with β-Galactosidase staining solution. Stained samples were visualized using a bright field microscope (Leica, Wetzlar, Germany).

### Proliferation assay

Cell proliferation rate was measured with a BrdU Cell Proliferation ELISA Kit (Abcam), according to the manufacturer’s instructions. Absorbance was monitored with a GloMax-Multi Detection System: Luminometer (Promega, USA).

### Wound healing assay

A reproducible wound was performed with a pipette tip on a confluent monolayer of WT, HGPS, HGPS-like and MAD-B fibroblasts cultured on 96-well plates. The medium was removed, and cells were incubated for 6 h with medium containing 500 nM MG132 or equal volume DMSO. The surface of the wound was acquired with a Zeiss Axio Observer using Zen 2.3 pro-software and measured with ImageJ software v1.52K. Results were expressed as a percentage of the area of the original wound, and normalized to DMSO-treated cells, considered as 100%.

### Multi-analyte ELISA array

Multi-array ELISA was performed using the Multi-Analyte ELISArray Kit (Qiagen, Hilden, Germany) according to the manufacturer’s instructions. In brief, the supernatants were centrifuged for 10 minutes at 1,000 × g to remove any particulate material. 50 μl of each experimental sample, was added to the array coated with specific cytokine capture antibodies: IL-1α IL-1β, IL-2, IL-4, IL-6, IL-8, IL-10, IL-17α, TNF-α, IFN-γ, TGFβ and GROa and incubated at room temperature (RT) for 2 hours. After three washes, 100 μl of the diluted biotinylated detection antibodies were added to the appropriate wells of the ELISA plate and incubated in the dark for 1 h at RT. The plate was washed three times and 100 μl of diluted Avidin-horseradish peroxidase (HRP) were added into all wells and incubated in the dark for 30 minutes at RT. Development and stop solutions were added followed by detection of absorbance at 450 on a Luminometer: GloMax-Multi Detection System (Promega, USA).

### Statistics

Statistical analyses were performed with the GraphPad Prism software. Differences between groups were assayed using a two-tailed Student’s t-test. In all cases, the experimental data were assumed to fulfill t-test requirements (normal distribution and similar variance); in those cases, where the assumption of the t-test was not valid, a nonparametric statistical method was used (Mann–Whitney test). A P-value less than 0.05 was considered as significant. Error bars indicate the standard error of the mean.

## ACKNOWLEDGEMENTS

We warmly acknowledge all family members and their relatives for their participation to this study. This work was supported by the Association Française contre les Myopathies (AFM) (AFM grant MNH-Decrypt 2011-2015 and TRIM-RD 2016-2020 to A.D.S.G. and N.L.), the Institut National de la Santé et de la Recherche Médicale (INSERM) (recurrent grants) and Aix-Marseille University (AMU).

## CONFLICTS OF INTEREST

The authors declare no competing or conflicting interests.

## AUTHOR CONTRIBUTIONS

KH, ADSG, and NL conceived and designed the experiments. KH performed the experiments and analyzed the data under the supervision of NL and ADSG. KH, NL, ADSG and PC wrote the manuscript. FC, GKM, YD, LVM and GMB carried out the clinical characterization and follow up of the patients. All authors read and approved the final manuscript.

## DATA AVAILABILITY

The authors state that all data generated during this study are included in the article, and that they are available from the corresponding author upon reasonable request. RNA seq data that support the findings of this study have been deposited in ArrayExpress under the accession numbers: E-MTAB-5807 (Harhouri; 2017-06-1; RNA seq of HGPS treated cells; ArrayExpress; E-MTAB-5807).

## SUPPORTING INFORMATIONS-FIGURES

**Figure S1.**
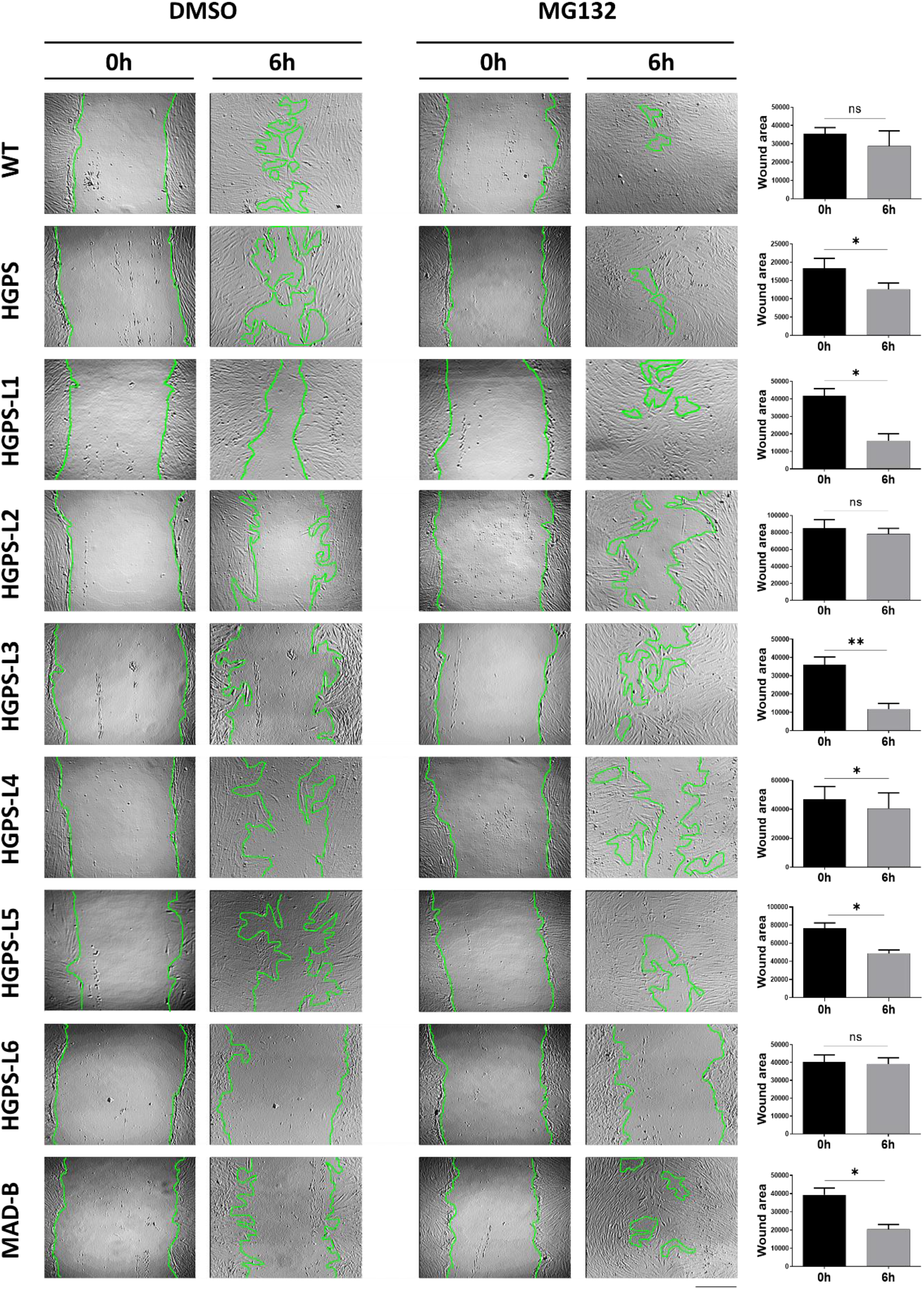
MG132 promotes HGPS-like and MAD-B fibroblasts migration. Wound healing assay performed on WT, HGPS, HGPS-like and MAD-B fibroblasts treated for 6h with DMSO or MG132 (500 nM). After subtracting the DMSO-induced wound repair during 6 h (control), results were expressed as the wound area following MG132 treatment (6 h) compared to the original wound (0 h). (mean ± SEM, n = 3, Student’s t-test, * p < 0.05, ** p < 0.01). Scale bar, 100 μm.

**Figure S2.**
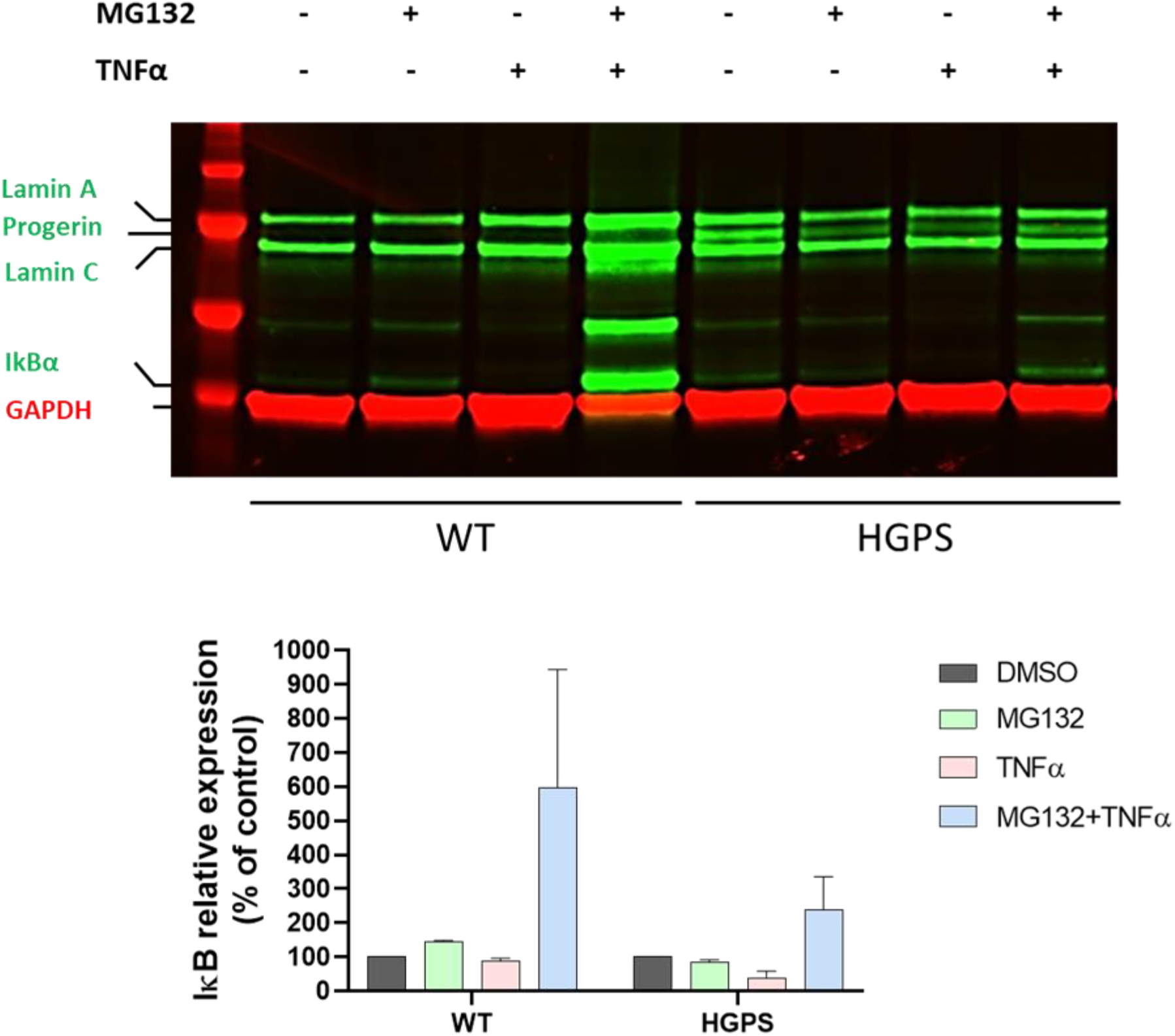
MG132 blocks the degradation of NF-κB inhibitor, I-κB. Upper panel: Western blotting evaluation of Lamin A/C and I-κB in whole lysates from WT and HGPS fibroblasts treated with DMSO (-), 500 nM MG132 for 48 h (+), 10 ng/ml TNFα for 48 h (+) alone or in combination. Lower panel: I-κB expression levels were normalized to GAPDH values using ImageJ software.

**Figure S3.**
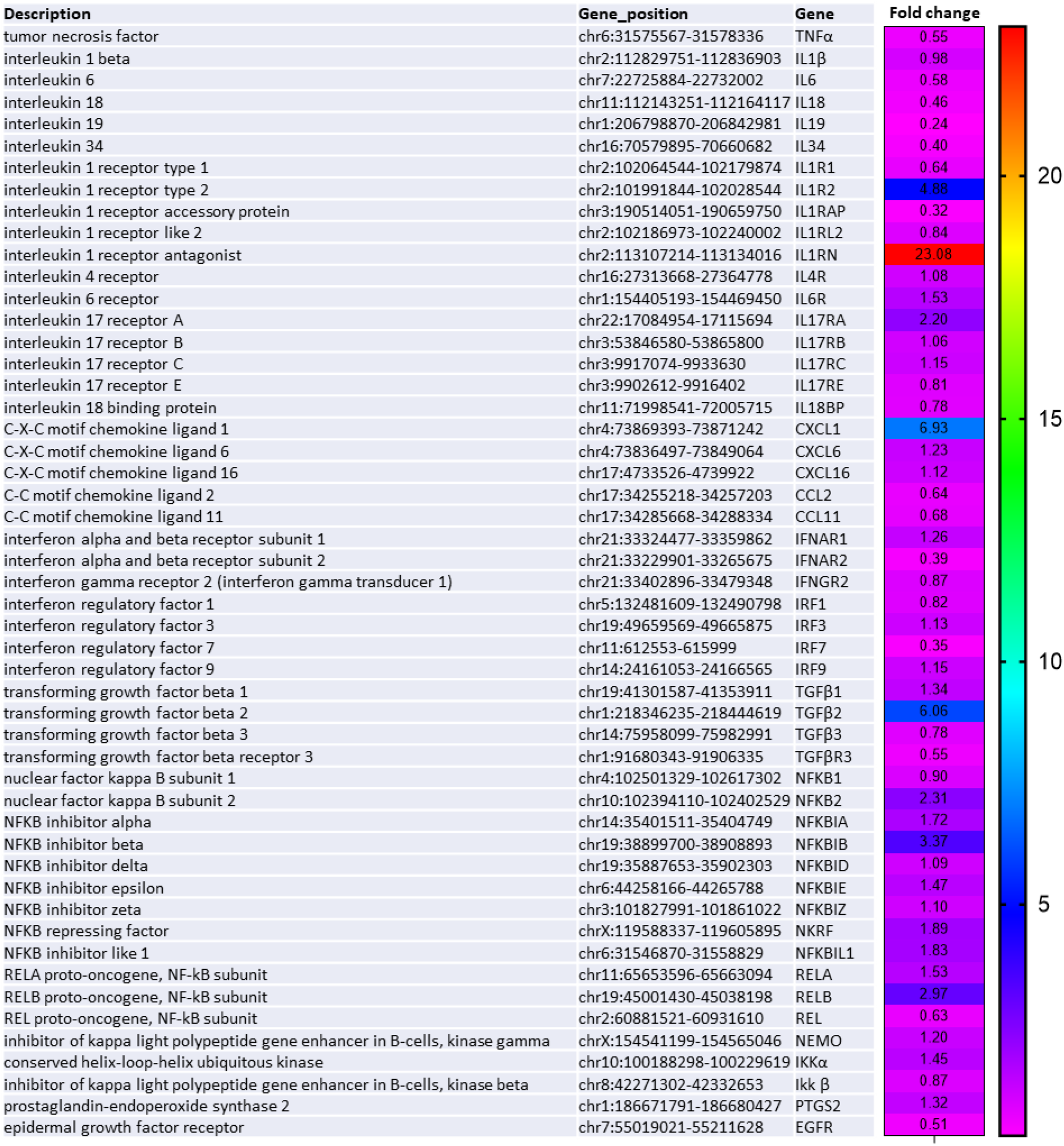
Transcriptional attenuation of inflammatory response to MG132 in classical HGPS fibroblasts. Heatmap of RNAseq data (ArrayExpress accession number: E-MTAB-5807) from HGPS fibroblasts treated with DMSO (vehicle control) or with 5 μM MG132 for 6 h. This analysis represents the fold change, in MG132-treated relative to DMSO-treated HGPS fibroblasts, of the most characteristic transcripts of the NF-κB pathway. (n = 2).

**Figure S4.**
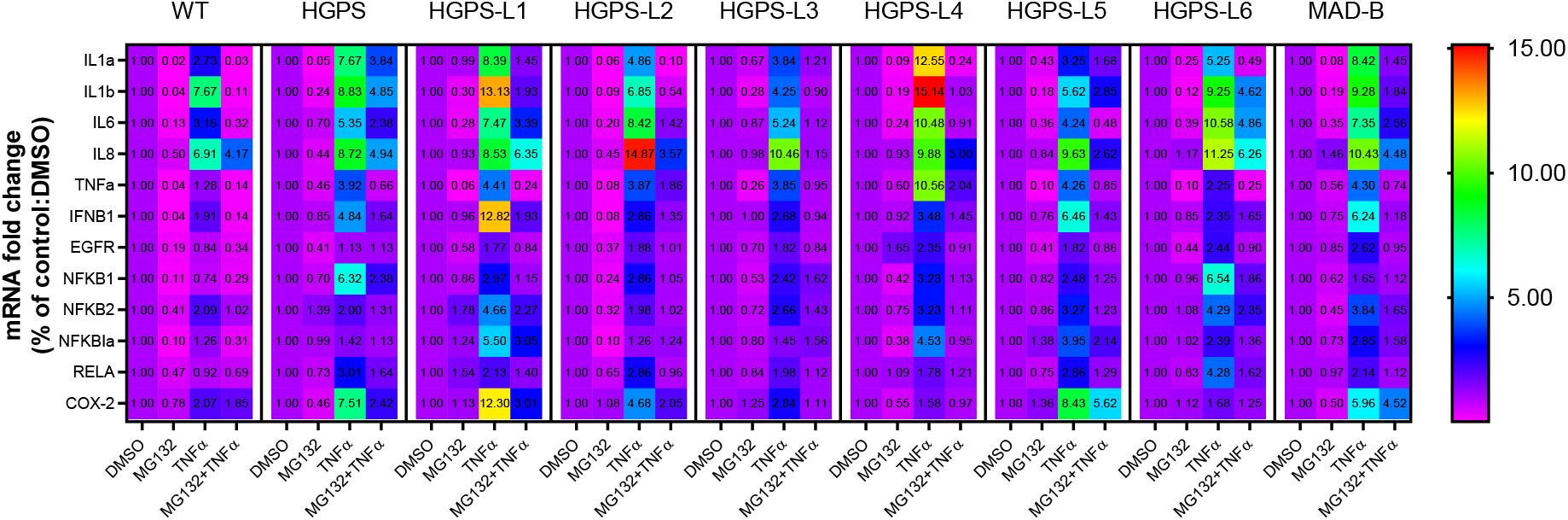
MG132 reduces the transcript levels of proinflammatory mediators and counteracts TNFα-induced inflammation. Quantitative real-time PCR using selected inflammatory genes expression arrays in culture supernatants of WT, HGPS, HGPS-like and MAD-B fibroblasts treated for 6 h with MG132 (500 nM), TNFα (10 ng/ml) alone and in combination or DMSO as a vehicle control.

**Figure S5.**
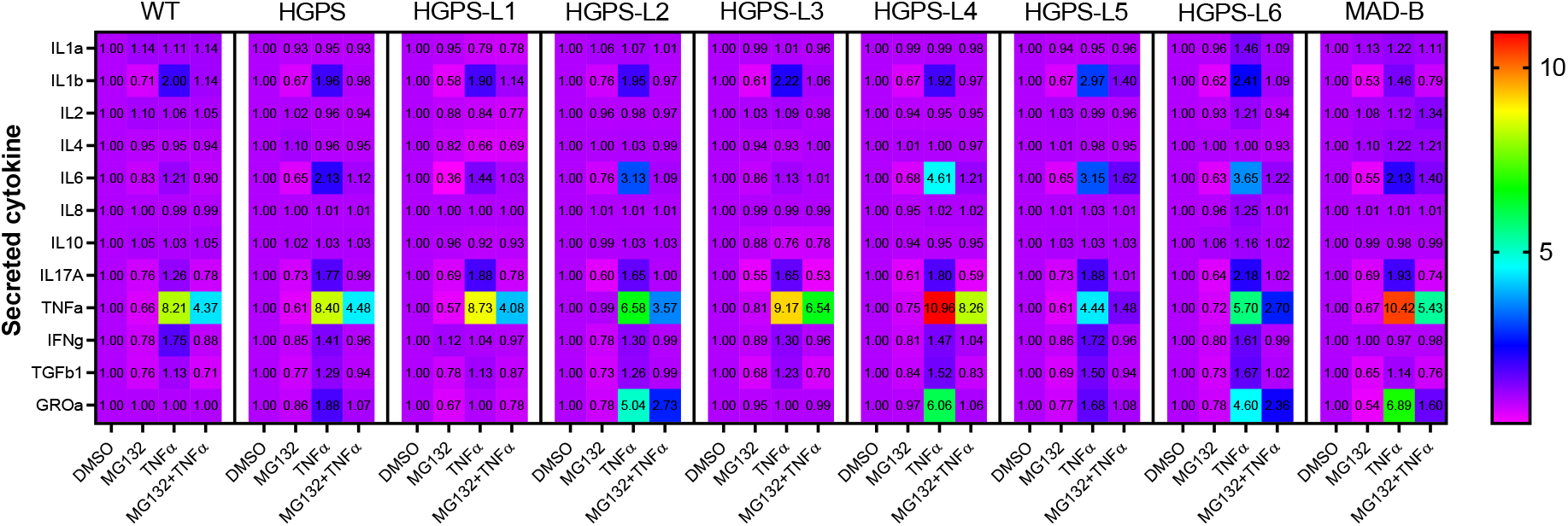
MG132 reduces the secretion of proinflammatory cytokines and alleviates TNFα-induced inflammation. Enzyme-Linked Immunosorbent Assay (ELISA) using multi-analyte ELISA arrays to measure inflammatory cytokines in culture supernatants of WT, HGPS, HGPS-like and MAD-B fibroblasts treated for 24 h with MG132 (500 nM), TNFα (10 ng/ml) alone and in combination or DMSO as a vehicle control.

## Notes

### Competing Interest Statement

The authors have declared no competing interest.

